# The role of innate immunity, antibiotics, and bacteriophages in the course of bacterial infections and their treatment

**DOI:** 10.1101/2024.08.23.609294

**Authors:** Brandon A. Berryhill, Teresa Gil-Gil, Christopher Witzany, David A. Goldberg, Nic M. Vega, Roland R. Regoes, Bruce R. Levin

**Author notes:** Corresponding Author: Bruce R. Levin, **Email:**.

## Abstract

Critical to our understanding of infections and their treatment is the role the innate immune system plays in controlling bacterial pathogens. Nevertheless, many in vivo systems are made or modified such that they do not have an innate immune response. Use of these systems denies the opportunity to examine the synergy between the immune system and antimicrobial agents. In this study we demonstrate that the larva of *Galleria mellonella* is an effective in vivo model for the study of the population and evolutionary biology of bacterial infections and their treatment. To do this we test three hypotheses concerning the role of the innate immune system during infection. We show: i) sufficiently high densities of bacteria are capable of saturating the innate immune system, ii) bacteriostatic drugs and bacteriophages are as effective as bactericidal antibiotics in preventing mortality and controlling bacterial densities, and iii) minority populations of bacteria resistant to a treating antibiotic will not ascend. Using a highly virulent strain of *Staphylococcus aureus* and a mathematical computer-simulation model, we further explore how the dynamics of the infection within the short term determine the ultimate infection outcome. We find that immune activation in response to high densities of bacteria leads to a strong but short-lived immune response which ultimately results in a high degree of mortality. Overall, our findings illustrate the utility of the *G. mellonella* model system in conjunction with established in vivo models in studying infectious disease progression and treatment.

**Significance statement:** Central to our understanding of the course of bacterial infections and their treatment is the contribution of the innate immune system. We use the larvae of *Galleria mellonella* to test hypothesis about the role of the innate immune system on *Staphylococcus aureus* infections. We demonstrate that the innate immune system of these larvae can control the infection and be saturated by high bacterial densities. As a consequence of this innate immune system, bacteriostatic drugs and phages are as effective as bactericidal drugs, and minority populations of bacteria resistant to antibiotics do not ascend. Our findings illustrate the utility of *G. mellonella* as a model for studying infections dynamics and therapeutic strategies.

## Introduction

The design of regimens for antibiotic therapy are based almost exclusively on studies of the pharmacodynamics of combinations of bacteria and antibiotics (1). These studies nearly always neglect the role of the host immune system in controlling infections, and thereby the synergy between antibiotics and the innate immune system to the course of treatment (2, 3). It is well known that an infected host’s immune system plays a major role in the clearance of bacterial infections (4). The omission of the immune response also applies to studies with animal models that are specifically designed and intended to increase our knowledge of infections beyond what we learn from in vitro experiments. These animal model studies commonly rely on mice that have no innate immune system (5-8). Prior to infection and treatment these mice are treated with agents such as cyclophosphamide, thereby making them neutropenic (9). Consequently, studies with these mice do not further our understand of the contribution of the innate immune system to the course of treatment of infections with antimicrobial agents.

Theoretical studies suggest and a few experiments with non-immunocompromised animals have generated the following hypotheses: i) in the absence of chemotherapy, if the density of the infecting bacterial population is great enough to saturate the innate immune system, the infection will fail to be controlled (10); ii) bacteriostatic antibiotics and bacteriophages can be as effective as bactericidal drugs in controlling the infection (10, 11); and iii) minority populations of bacteria that are resistant to the treating antibiotic will not ascend (12). These results are predicated on the function of the innate immune system. Innate immunity is the primary factor controlling bacterial density (hypothesis i) irrespective of the mechanism of treatment (hypothesis ii) and is responsible for the suppression of infrequent bacterial populations (hypothesis iii).

To test these hypotheses and explore the role of the innate immune system in the control of bacterial infections and test the above hypotheses, we employ, in a quantitative fashion, the larvae of the Wax Moth, *Galleria mellonella (13)*. These larvae have several qualities which make them amenable as a model system for this study: i) they have an innate immune system (14), primarily hemocytes (which are analogous to the neutrophils of mammals) and melanocytes; ii) the morbidity and mortality of these larvae is reflected by a change in melanization—which is to say a darkening of the larva, serving as a useful early indicator (15); iii) they are inexpensive and experiments can be highly replicated; iv) manipulation of this system is relatively facile and has no bioethical constraints; v) the densities of infecting bacteria can readily be estimated; vi) there is an extensive literature of using this model to discover new anti-infective compounds, particularly anti-fungals (16); vii) several studies of drug kinetics have been performed in this system (17, 18); and viii) there are survival studies addressing the role of virulence and treatment efficacy (19, 20).

In this study we first confirm that those results obtained with *G. mellonella* mirror those obtained in other systems employed for studying the dynamics of infection, particularly those with non-neutropenic mice. Using these larvae and *Staphylococcus aureus*, we explore the appropriateness of this system for testing the above hypotheses. Lastly, we quantitively characterize the dynamics of bacterial infection with a highly toxigenic strain of *S. aureus* (21). To explore an unanticipated result obtained from the dynamics of infection, we employ a mathematical computer-simulation model.

## Results

### Larvae Survival Is Dependent on the Density of Infecting Bacteria

Previous work in mice has shown that mortality is dependent on the initial bacterial inoculum (22). Critically, there appears to be a threshold effect under which mortality is mild, but once the inoculum exceeds that threshold density, mortality rises sharply. This result also obtains in *G. mellonella* when they are infected with *S. aureus*, where we find the critical threshold to be approximately 10^6^ CFU/larva (Figure 1). Our experiments support the hypothesis that in the absence of chemotherapy, if the density of the infecting bacterial population is great enough to saturate the innate immune system, the infection will fail to be controlled.

**Figure 1.**
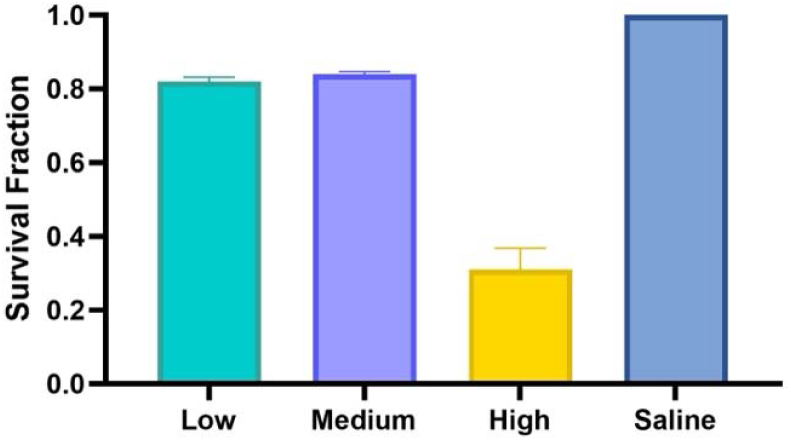
Larvae survival at 24 hours with different bacterial inoculum densities. Larvae were injected with *S. aureus* MN8 at 10^4^ CFU/larva (turquoise), 10^6^ CFU/larva (purple), 10^8^ CFU/larva (yellow), or saline (blue). Mortality was assessed at 24 hours. Presented are means and standard deviations for the fraction of survival (N= 80 larvae). Low, medium, and high differ with saline at *p*<.0001^****;^ low is statistically different from high at *p*<.001 ^***;^ medium is different with high at *p*<.001 ^***;^ and low and medium are not statistically different.

### Treatment With Antibiotics and Phage Increases Larvae Survival

Theory predicts, and limited experiments confirm, that treatment of an infection with antibiotics will increase survival and that this outcome is independent of the type of antibiotic (23). To test these predictions in our *in vivo* system, we infected larvae of *G. mellonella* with a high density of *S. aureus* and treated immediately. All therapies tested increased survival to 90-100%, supporting the idea that bacteriostatic and bactericidal antibiotics are equally effective when the innate immune system is present (Figure 2A). Interestingly, treatment with a highly lytic bacteriophage, PYO^Sa^ (24), was able to replicate in these larvae (Supplemental Figure 1) and subsequently increase survival of the larvae by 10-fold, a result comparable to the antibiotics used.

**Figure 2.**
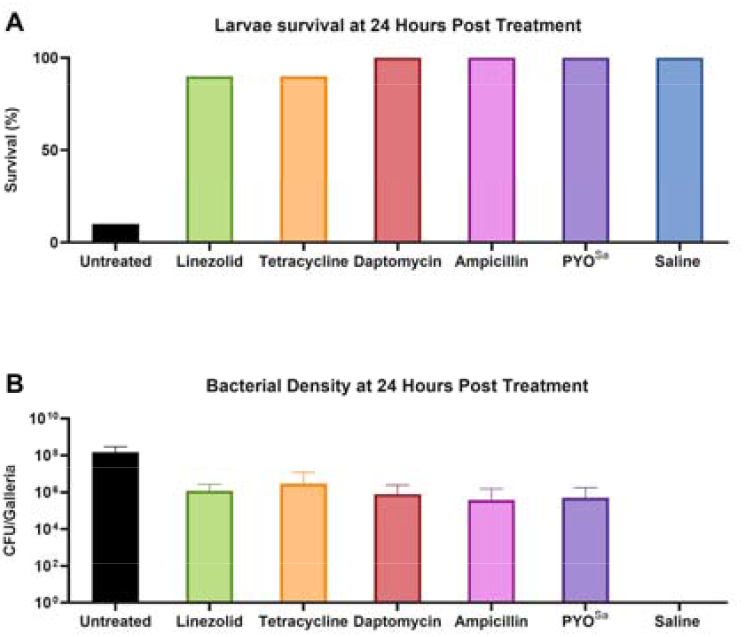
The effects of treatment on survival and bacterial density. Larvae were injected with 10^8^ CFU/larva of *S. aureus* MN8 and either were untreated (black) or immediately treated with linezolid (green; a bacteriostatic drug), tetracycline (orange; a bacteriostatic drug), daptomycin (red; a bactericidal drug), ampicillin (pink; a bactericidal drug), or the phage PYO^Sa^ (purple). 24 hours after treatment, (A) mortality was assessed and compared to an uninfected saline injection control (blue) and (B) the final bacterial density was determined. Shown are means and standard deviations (N=20 larvae). All treatment conditions statistically differ from the saline in terms of survival and density (*p*<.0001 ^****^ and *p*<.0001 ^****^, respectively).

All therapies, be they bacteriostatic antibiotics, bactericidal antibiotics, or lytic phage, decrease bacterial load to roughly the same level. While treatment drastically increases survival, it does not clear the infecting bacteria: however, all treatments decrease bacterial densities to around or below the critical threshold of 10^6^ CFU/larva. This result is consistent with immune exhaustion being responsible for mortality during a bacterial infection. Of note, the strain of *S. aureus* used here, MN8, produces a beta-lactamase and has an ampicillin minimum inhibitory concentration in excess of 256 μg/mL, demonstrating that the effective treatment can occur even when the underlying bacteria is resistant to the treating antibiotic.

### Antibiotic-Resistant Bacteria do not Increase in Frequency Under Antibiotic Treatment

It has been postulated from mathematical modelling (25) and demonstrated using mice (12) that one role the innate immune system plays in controlling infections is preventing the increase in frequency of rare antibiotic resistant mutants. To explore if this result would obtain with *G. mellonella*, larvae were infected with rifampin resistant and rifampin sensitive *S. aureus* MN8 at several ratios ranging from 0.001 to 1 up to 1 to 1. Infected larvae were immediately treated with rifampin and the ratio of resistant to sensitive cells enumerated after 24 hours (Figure 3). As anticipated, the antibiotic-resistant population did not substantially increase in the presence of antibiotic treatment (Figure 3A). This result, however, does not obtain when the immune system of the larva is ablated (Figure 3B). With an ablated immune system, the minor population of antibiotic-resistant bacteria increases in frequency much like in the in vitro condition (Figure 3C).

**Figure 3.**
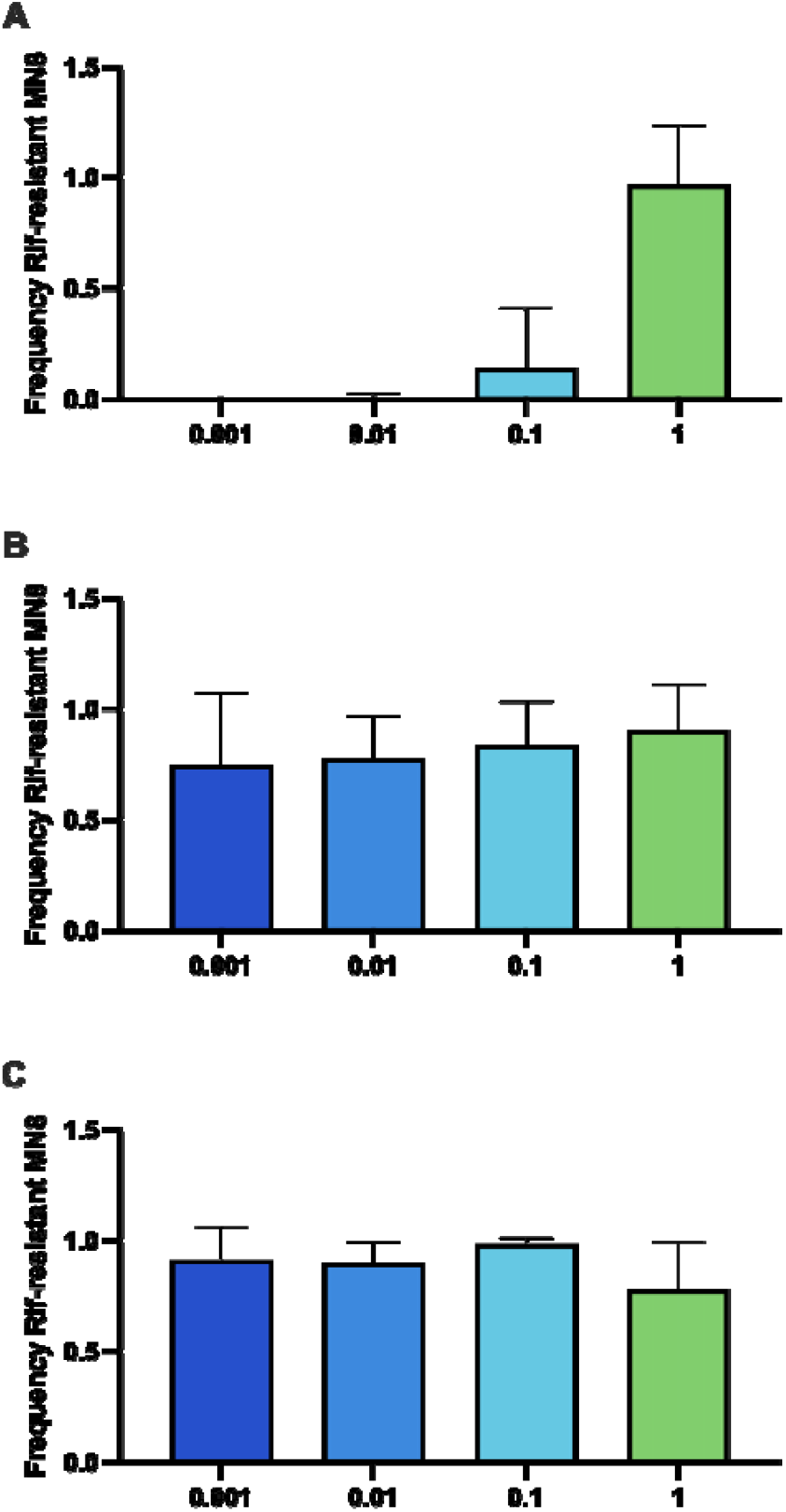
Invasion when rare of an antibiotic-resistant subpopulation. (A) In vivo with a functional immune system; (B) in vivo with the immune system of the larvae knocked out; (C) in vitro. At ratios of 0.001, 0.01, 0.1, or 1 to 1, rifampin-resistant *S. aureus* MN8 and rifampin-susceptible *S. aureus* MN8 were injected into 20 larvae with either a functional or ablated immune system then immediately treated with rifampin. After 24 hours, larvae were sacrificed and the final ratio of rifampin resistant to rifampin sensitive bacteria determined. As a control for the in vivo invasion when rare, at the same ratios, rifampin-resistant *S. aureus* MN8 and rifampin-susceptible *S. aureus* MN8 were mixed and immediately treated with rifampin in vitro. Flasks were grown overnight and the ratio of rifampin resistant to rifampin sensitive cells in vitro determined. Presented are means and standard deviations at 24 hours. The difference in the final ratios at all inoculum conditions condition except 1:1 are statistically different between panels A and B and A and C (*p*<0.0001 ^****^), however there is no significant difference between panels B and C.

### The Innate Immune System Functions via ETosis

*G. mellonella* have three components to their innate immune response (26): i) antimicrobial peptides (AMP) and lytic enzymes, ii) melanocytes, and iii) hemocytes. Since *S. aureus* MN8 has many anti-AMP responses, this humoral response does not contribute to the dynamics of infection. Further, since the infections are systemic, the production of melanin by melanocytes is not able to sequester the *S. aureus*, thus this response plays little role in controlling the infection (14). This leaves only hemocytes as the main determinant of the immune response. These hemocytes, upon detection of lipopolysaccharides or wall teichoic acid, release their DNA which forms an extracellular trap (ET) in a process called ETosis (27). Here, we show that the presence of a GFP labeled *S. aureus* is sufficient to cause ETosis (Figure 4A) and that these ETs can be degraded by DNase (Figure 4B).

**Figure 4.**
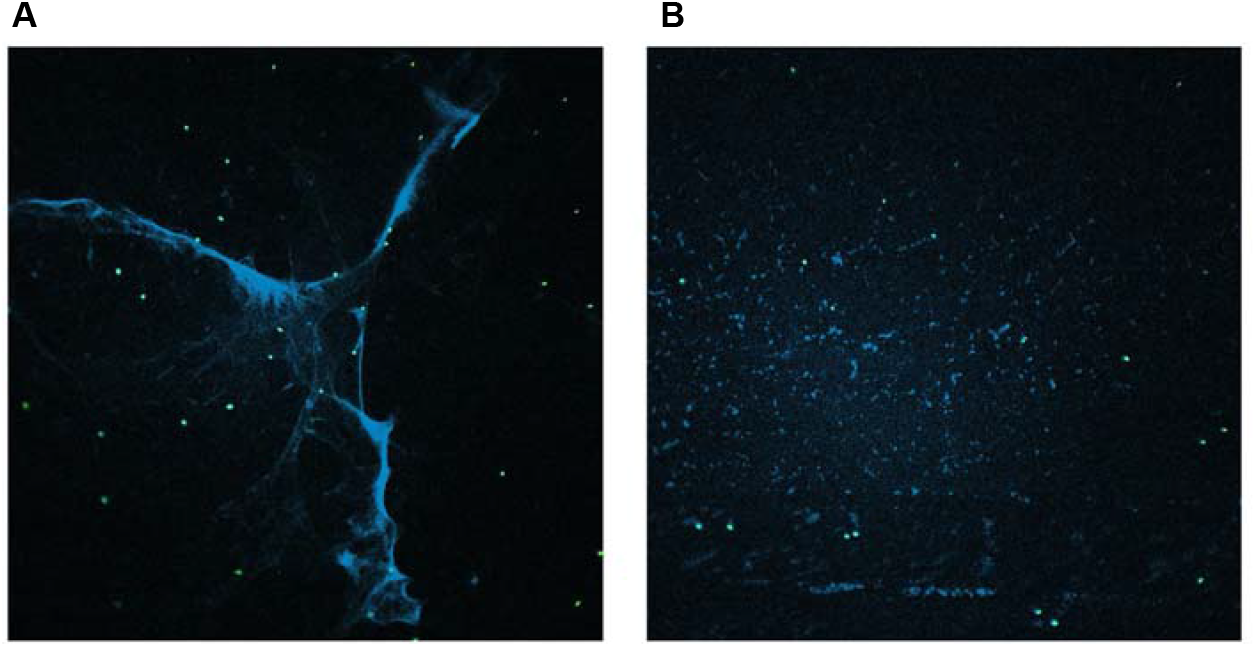
Hemocytes producing ETs. Hemocytes were incubated for 30 minutes with GFP producing *S. aureus* before being treated with either saline (A) or DNAse (B) and then stained with DAPI.

### Dynamics of Infection in *G. mellonella*

To understand the dynamics of how the immune system controls infections with *S. aureus* of differing initial densities, we infected 160 larvae with 10^4^, 10^6^, or 10^8^ CFU/larva and sacrificed half of each condition at 4 hours (Figure 5A) and the other half at 24 hours (Figure 5B). Shown in Supplemental Figure 2 are the dynamics of each timepoint, and inoculum density separated. Of note, at 4 hours there is a convergence of all initial densities to around 10^5^-10^6^ (the critical density), illustrating that the immune system can control the infection in the short term regardless of initial infection density. However, at 24 hours, while the 10^4^ CFU/larva and 10^6^ CFU/larva initial inoculum infections are still controlled and around the same density as at 4 hours, the 10^8^ CFU/larva initial inoculum infections have dramatically increased in density, corresponding with the high mortality seen in Figure 1. We hypothesize that the reason for the failure of the immune system to control the infection at 24 hours while it was readily capable of doing so at 4 hours is immune exhaustion. The hemocytes release their DNA in a density-dependent manner,thus, to control the high inoculum infection at 4 hours, a large number of the hemocytes had to release their DNA and ultimately die, leading to an exhaustion of these immune cells, leading to the infection not being controlled by 24 hours. To ensure that the results observed at 24 hours are the endpoints of the infection, rather than an intermediate stage, larvae were infected and maintained for 72 hours (Supplemental Figure 3). Notably, both survival and infection dynamics are qualitatively similar to that observed at 24 hours, with the exception that the average density of bacteria per larvae in the low inoculum group is several logs lower at 72 hours—showing a higher degree of infection control.

**Figure 5.**
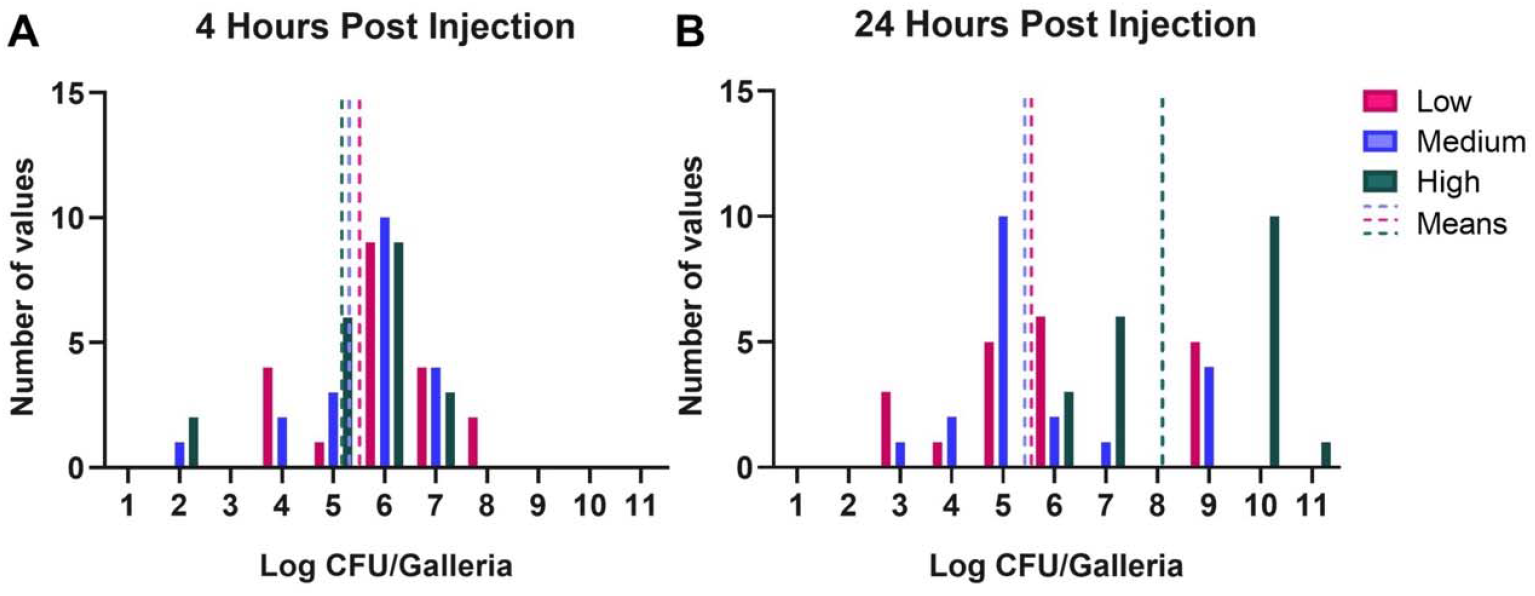
Dynamics of infection. Larva were infected with three different densities of *S. aureus* MN8: low (10^4^ CFU/larva; pink), medium (10^6^ CFU/larva; purple), or high (10^8^ CFU/larva; green) and were sacrificed after (A) 4 or (B) 24 hours post infection. Shown are the number of larvae within each log_10_ CFU. The mean of the CFU at each timepoint for each inoculum density is presented as a dotted line of the same color. For each condition N=20 individual larvae.

### A Model of the Observed Infection Dynamics

We developed a mathematical population model that incorporates the innate immune system to explain the within-host dynamics of bacteria observed in the infected larva. By explicitly modeling both the bacterial population and the host immune response, we can evaluate how different initial bacterial population sizes affect infection dynamics and assess when the immune system is overwhelmed. We simulate the innate immuneresponse under the assumption that immune effectors (*E*) need to be activated, then encounter (unprotected) bacteria (*U*), and subsequently take time to kill them (28). Further, our model accounts for persistent infections with low population sizes (as observed in Figure 2B) by incorporating a protected sanctuary site where bacteria evade the immune system (*P*). See Supplemental Information for model equations and detailed description.

Our simulations show that infection dynamics highly depend on the initial size of the inoculum (i.e., the starting population of bacteria *U*_O_). For simulations with inoculum sizes above 10^5^ cells, the total bacteria population grows to around 10^8^ cells within 15 hours (Figure 6). This growth is mainly driven by the unprotected bacteria *U* and limited by the carrying capacity of the unprotected compartment (*K*_*U*_, Table S1). Over the same time frame, the activated immune effectors *E* decline and ultimately reach zero. This decline shows that all *E* are engaged with bacteria *U*, but still cannot kill *U* faster than they are growing, hence the infection cannot be controlled, and the immune system gets overwhelmed.

**Figure 6.**
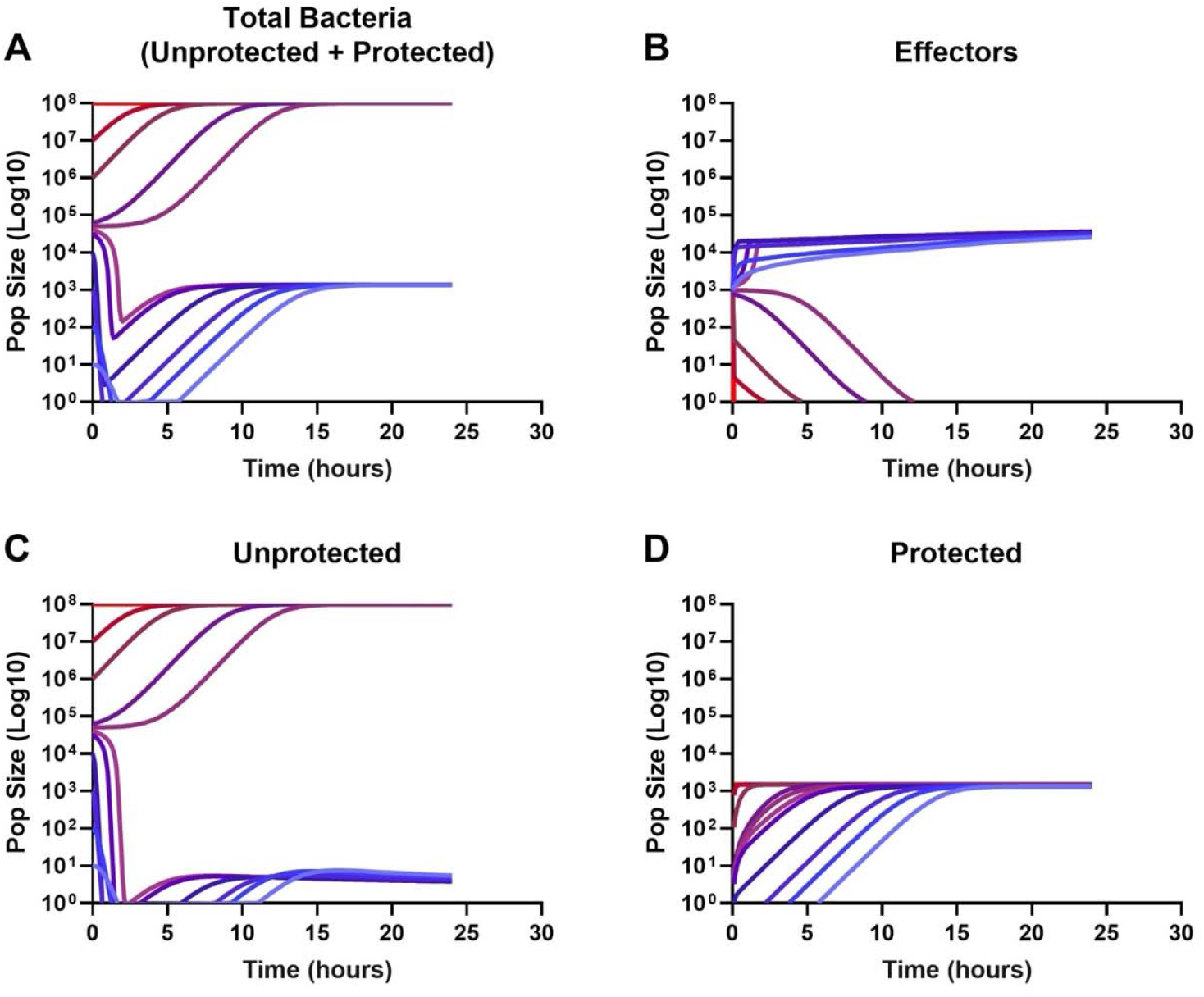
Simulated dynamics of within-host bacterial infections. Shown are the population sizes over time of (a) the total bacteria present within the host (sum of *U* and *P*), (b) the number of activated host immune effectors (*E*), (c) the number of bacteria in the unprotected site (*U*) and (d) the protected site (*P*). Each panel shows multiple-colored lines, with colors representing individual simulation runs with increasing inoculum sizes (blue to red). The following parameter values were used for this simulation: r = 1, K_u_ = 10^8^, K_p_ = 1.5×10^3^, f = 0.001, b = 0.1, E_0_ = 10^3^, E_tot_ = 10^5^, h_1_ = 0.001, h_2_ = 0.001, d = 0.01, a = 0.01, g = 0.5, and s = 0.001. See Methods and Table S1 for model and parameter descriptions.

For inoculum sizes below 10^5^ we observe an initial drop in the total bacterial population, caused by the removal of *U* by *E*. Simultaneously, in response to the presence of *U*, more effectors get activated resulting in an increase in *E* until the total of available effectors is reached (*E* _*tot*_ = 10^5^). The larger the initial population *U* the faster this activation happens. However, all simulations with inocula <10^5^ quickly start to regrow after this initial decline. This recovery is not driven by *U*, but rather by growth of *P* in the protected compartment. A fraction of bacteria *U* migrates from the unprotected to the protected compartment before being removed by the immune system, where they grow protected from *E* until they reach capacity (*K*_*p*_ = 1500) – this occurs faster with larger initial inocula. Consequently, the infections become controlled and persistent, because *E* can keep in check the bacteria which migrate out of protection, stopping the total bacterial population from growing further, but *E* are unable to eradicate the bacteria *P* in the protected compartment.

In summary, our simulations show that there is a critical threshold inoculum size of around ∼10^5^ cells, at or above which infections quickly overwhelm the immune effectors, i.e. the immune effector population goes to zero, and bacteria grow to capacity in both the unprotected (*U*) as well as the protected compartment (*P*) (Figure S4). If the inoculum is below this critical threshold the infection can be controlled by the immune effectors, and the final total population size corresponds to the carrying capacity of the protected site (*K*_*P*_). This critical threshold for the inoculum size, above which all infections become uncontrollable, exists for a wide range of parameters – and is primarily influenced by the immune effector killing rate (*h*_l_) and effector engagement rate (*h*_2_) (see Figure S5 and S6).

## Discussion

Critical to understanding both the course of an infection and how treatment changes that course is the innate immune system. Most in vivo systems traditionally used to study the treatment of bacterial infections purposefully ignore the immune response (5-8). In this study, we present a promising system that has an innate immune system which has elements which are analogous to those in humans, in that it works via ETosis, as neutrophils do. This highly tractable, inexpensive system with no animal biosafety considerations is the larva of the wax moth, *Galleria mellonella*. This system enabled us to test three major hypotheses about the functioning of the innate immune system during the course of an infection: i) in the absence of therapy, if the infecting bacteria is of sufficient density, the immune system will saturate and the infection will progress uncontrolled; ii) bacteriostatic antibiotics and phages will reduce mortality to the same degree as bactericidal drugs; and iii) minority antibiotic-resistant populations will not increase in frequency during antibiotic treatment.

While testing the above hypotheses with *G. mellonella*, we found support for each hypothesis that mirrors the previous results obtained in parallel experiments in mice. First, we demonstrated that the mortality is highly dependent on the initial inoculum such that initial densities above 10^6^ CFU per larva results in mortality above 85%. Second, we found evidence that both bacteriostatic antibiotics and phages were as successful as bactericidal drugs in reducing mortality as well as in reducing the bacterial density after 24 hours. Interestingly, ampicillin was as effective as the other treatments even though this strain of *S. aureus* is highly resistant to beta-lactams. This result, while unintuitive, mirrors that seen in clinics where it has been observed that if the underlying pathogen is resistant to the treating drug, treatment will still succeed 60% of the time (29, 30). Third, unlike in vitro experiments, when a larva is infected with a minority population of antibiotic-resistant bacteria, under treatment with the drug to which the minority population is resistant, this sub-population does not increase in frequency and invade when rare if the immune system is functional.

In addition to providing support to the above hypotheses, our results raise an interesting question about how the dynamics of infection underlie morbidity and mortality. When the initial inoculum is low (∼10^4^ CFU per larva) or medium (∼10^6^ CFU per larva), the infection density will rise slightly at 4 hours but be controlled at a fixed density over 24 hours. More importantly, survival will be nearly 100%. When the initial inoculum is high (∼10^8^ CFU per larva) the bacterial density will be brought down such that the density at 4 hours is the same as the low and medium inocula. In stark contrast to the other two conditions, by 24 hours, the density of bacteria in the larvae will not be controlled and often in excess of 10^8^ CFU per larva, leading to a mortality of nearly 90%.

These dynamics were unexpected and suggest that the immune system responds very aggressively when there is a high inoculum, such that the host’s cells responsible for the control of the infection are exhausted. This result is consistent with the mechanism by which these larvae control infections. When hemocytes detect lipopolysaccharides or (in this case) teichoic acids, they lyse and release their DNA in an effort to control the infection (31). Therefore, the higher the initial inoculum, the more immune cell death can be expected. Ultimately, the exhaustion of these cells leads to the failure of the immune system to control the infection by 24 hours. In support of this premise, we constructed amathematical computer-simulation model. With biologically plausible parameters and assumptions, the model was able to recapitulate the above-described experimental results and found a critical inoculum above which the immune system will not control the infection.

In supporting the use of *G. mellonella* more broadly as a model system for infections, one needs to be able to manipulate the system, in particular, their innate immune response. Others have performed genetic manipulation of the system as well as chemical ablation of the innate immune system (32-34). We also performed this chemical procedure on the larvae to determine the effects that removing the immune response has on the infection dynamics (Supplemental Figure 7). First, in support that the immune ablation was successful, we lose the dose-dependent response of morbidity, mortality, and infection dynamics. However, *a priori we* expected mortality to trend to 0 and the infection densities to be extremely high. This is the opposite of what we saw. Generally, survival was high and the bacterial density in the surviving larvae was extremely low (less than 10^1^ CFU/larvae in most cases). On the other hand, the density of bacteria in the dead larvae was high. This provides another line of evidence that 10^6^ CFU/larvae is this critical threshold for predicting survival. While we cannot currently experimentally explain these results, we do have a hypothesis we intend to test in the future. It is not currently clear where the *S. aureus* are located in the larva during an infection, however it is well known that Staphylococcus has an intra-phagocytic niche (35). As these larvae have an existing, stable microbiome, could it be that the only niche available for the infecting bacteria to fill is inside of these immune cells? Thus, when we ablate the immune response by killing the immune cells, the bacteria have nowhere to proliferate and thus are lost from the larvae. Supporting this hypothesis is the high density found in the dead larvae, in these cases due to technical error or some other uncontrolled variation, it is possible the chemicals were not able to fully ablate the immune system, thus leaving a niche for the bacteria to proliferate in, and now since the immune system is suppressed, the infection is fatal and the bacteria reach a high density.

While this study has focused on the dynamics of infection, future studies will seek to elucidate the conditions under which treatment with antibiotics and/or phages will be synergistic or antagonistic to the innate immune system dynamics observed here. This system, with its phenotypic marker for sickness, will enable us to treat infections not on a fixed timeline, but instead when the individual is displaying symptoms of morbidity. This allows a more realistic analogue to how humans are treated, as no one seeks medical attention when they become infected, but instead they wait until they are sick. Allowing this delay in treatment is vital to understanding the dynamics of infections under anti-infective therapy.

## Materials and Methods

### Growth media

All experiments were conducted in Muller Hinton II (MHII) Broth (90922-500G) obtained from Millipore. All bacterial quantification was done on Lysogeny Broth (LB) agar (244510) plates obtained from BD. E-tests were performed on MH agar plates made from MH broth (M391-500g) with 1.6% agar obtained from HiMedia. Overnights of GFP labeled S. aureus Newman were grown with 25 μg/mL of ampicillin.

### Growth and infection conditions

All experiments were conducted at 37 °C.

### Bacterial strains

All experiments were performed with *S. aureus* MN8 obtained from Tim Read of Emory University. *S. aureus* MN8 was marked with streptomycin resistance to enable differential plating from the larva microbiota. Green fluorescent protein labelled *S. aureus* Newman was obtained from Dr. Nic Vega of Emory University.

### Antibiotics

Daptomycin (D2446) was obtained from Sigma-Aldrich. Tetracycline (T17000) was obtained from Research Products International. Ampicillin (A9518-25G) was obtained from Sigma-Aldrich. Linezolid (A3605500-25g) was obtained from AmBeed. All E-test strips were obtained from Biomérieux.

### Bacteriophage

The bacteriophage PYO^Sa^ was obtained from the Levin Laboratory’s bacteriophage collection.

### Bacteriophage preparation for injection

Lysates of PYO^Sa^ were grown on *S. aureus* Newman such that the total volume exceeded 250 mL of media. These initial lysates were spun down and filtered through a 0.22 μm filter to remove cellular debris. From there, the lysates were run through a 100 kD tangent flow filtration (TFF) cassette (PAL 0A100C12) with a PAL Minimate TFF system. The volume was reduced during this process to 15 mL which was refiltered through a 0.22 μm filter. This lysate was incubated with shaking at 25 ^°^C with equal parts octanol (Thermo Scientific 4345810000) for 24 hours. The non-organic fraction was then dialyzed with Thermo Scientific’s 250 kD float-a-lyzer cassette (66455) against 60% ethanol and then again against saline. Finally, the lysate was refiltered through a 0.22 μm filter.

### Sampling bacterial and phage densitie

Bacteria and phage densities were estimated by serial dilutions in 0.85% NaCl solution followed by plating. The total density of bacteria was estimated on LB (1.6%) agar plates. To estimate the densities of free phage, chloroform was added to suspensions before serial dilutions. These suspensions were mixed with 0.1 mL of overnight MHII grown cultures of wild-type *S. aureus* MN8 in 4 mL of LB soft (0.65%) agar and poured onto semihard (1%) LB agar plates.

### *G. mellonella* preparation

*G. mellonella* larvae were obtained from Speedy Worm (Minnesota, USA) and placed immediately at 4 ^°^ C for 24 hours as a cold shock. Larvae were then sorted such that only those that weighed between 180 and 260 mg.

### *G. mellonella* infection and sampling

Larvae were allowed to acclimatize at 37 ^°^ C for 24 hours before the experiment. Overnight cultures of *S. aureus* MN8 were centrifuged for 20 min at room temperature at maximum speed, the supernatant discarded, and the pellet resuspended in saline. This process was repeated 5 times. The bacterial suspension was diluted in saline to the desired inoculum concentration. Groups of 10 larvae were injected with 5 μL of the desired bacterial suspension at the last left proleg. Larvae were incubated at 37 °C in the dark without food. After 4 or 24 hours the larvae were placed in 1 mL of saline and then homogenized. The homogenate was plated in LB agar plates containing 400 μg/mL of streptomycin.

### *G. mellonella* treatment

Larva were treated by injection in the last right proleg. Antibiotic concentrations were analogous to those used for clinical treatment in humans and the amount of antibiotic determined by weight for a total final treatment amount of: LIN-0.002 mg; TET-0.01 mg; DAP-0.002 mg; AMP-0.04 mg.

### *G. mellonella* immune system ablation

As in (32), larvae were injected in the right proleg with 20 μL of 10 μM cytochalasin b and nocodazole (note: this protocol does not work with 10 μL of 20 μM of each drug) and allowed to incubate in the dark at 37 °C for 4 hours. At 4 hours, larvae were infected as described above with a suspension that is 50% bacteria and 50% of the mixture of both of these drugs.

### Antibiotic minimum inhibitory concentration

Resistance to ampicillin was determined by E-test on MHII agar plates.

### Hemolymph extraction and hemocyte isolation

Larvae were dipped in 100% ethanol and allowed to dry. After drying, larvae were stabbed in a proleg once an hour for four hours with a 30-gauge needle (BD, 305195) to stimulate the innate immune system (36). After the fourth timepoint, the larvae were stabbed in the central line and the hemolymph removed and placed in 1 mL of Insect Physiological Saline (IPS; 150mM Sodium Chloride, Fisher, S271; 5mM potassium chloride, Sigma, P-8041; 100mM Tris/HCl, Fisher, BP153; 10mM EDTA, Promega, V4233; 30 mM sodium citrate, J.T. Baker, 3650) The lymph from 10-15 worms were pooled, the Eppendorf tube was centrifuged at 500 x g for 10 minutes. The pellet was washed twice with cold IPS then resuspended in a minimal amount of IPS.

### Microscopy

GFP-marked *S. aureus* was incubated with 10^6^-10^7^ hemocytes for 30 minutes in an Eppendorf tube. After incubation with bacteria, 10 µL of the cells were stained with 10 µL of 0.4% trypan blue (Sigma, T8154), mounted and imaged on a Leica DMi8 motorized inverted microscope with motorized stage (Leica, 11889113) to ensure cells were viable. After ensuring hemocytes are viable, the sample was divided in half. Half of the sample was treated with PBS and the other half of DNAse (MilliporeSigma, AMPD1) both samples were fixed with 4% paraformaldehyde (ThermoFisher, 043368.9M) and stained with 2-(4amidinophenyl)-1H-indole-6-carboxamidine (DAPI, 1 µg/mL; Invitrogen, D1306) and washed with Dulbecco’s PBS. Afterwards, 2 drops of antifade mountant (Invitrogen, P36980) was applied and slides allowed to cure on benchtop for 24 hours. The cells were then imaged on a Nikon A1R HD25 confocal microscope system at 60x oil immersion lens with NA = 1.49.

### Statistical analysis

All statistical analyses were performed in GraphPad Prism Version 8.0.1 using a two-tailed unpaired parametric t-test.

### Mathematical Model

We model infection and immune response dynamics by the following ordinary differential equations:

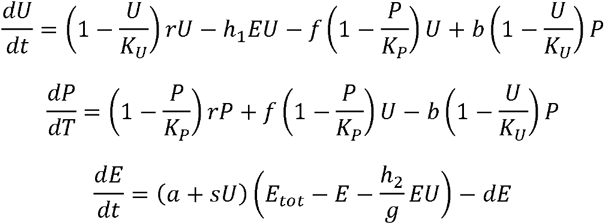

*U* and *P* describe bacterial populations in compartments where they are either unprotected or protected from the active immune effector population *E*. Both, *U* and *P* grow at rate *r*, limited by the carrying capacities *K*_*U*_ and *K*_*P*_, respectively. Bacteria migrate from *U* to *P* at rate *f* and from *U* to *P* at rate *b*, limited by the free capacity in the destination compartment, i.e.,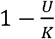. Immune effectors *E* are activated from a limited pool *E*_*tot*_ at background activation rate *a*, or in response to bacteria *sU*, and deactivate at rate. Unprotected bacteria *U* are removed by rate *h*_l_ proportional to the product of *U* and *E* (*h*_l_*UE*). Immune action is modelled as a two-step process: engagement at rate *h*_2_ (akin to capture) and killing at rate *g* (akin to digestion). Assuming *g* is large, the total number of un-activated immune effectors can be described by 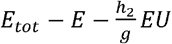 where 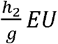 reflects effectors currently engaged with *U* (28). See Supplemental Text for details.

### Numerical solutions (simulations

All numerical simulations were implemented in R (version 4.3.1) using the deSolve package (37). Parameter sampling for the sensitivity analysis was done using Latin Hypercube sampling via the lhs package to efficiently maximize coverage of the parameter space (38). Global parameter sensitivity analysis was performed using the PAWN method implemented in the SAFER package (39). Briefly, the PAWN method evaluates parameter importance by the assessing the difference in output distributions if all parameters can vary compared to when one parameter is fixed. See Text S1 for equations and detailed description of the mathematical model, and Table S1 for parameter descriptions, values, and sampling ranges of the sensitivity analysis. All simulation code is available under https://gitfront.io/r/user-2939733/ruxwXHz8cVY9/galleria_immune/.

## Supporting information

All Supplemental Materials

## Data Availability

All data underlying this report can be found in the main text and its figures or the supplemental materials. Access to bacterial strains and phages can be had by contacting the corresponding author.

## Author Contributions

Conceptualization: BAB, BRL

Methodology: BAB, TGG, CW

Investigation: BAB, TGG, CW, DAG

Visualization: TGG

Funding Acquisition: BRL, RRR

Project Administration: BRL Supervision: BRL, RRR

Writing– Initial Draft: BAB, TGG, CW, BRL

Writing– Review & Editing: BAB, TGG, CW DAG, NMV, RRR, BRL

## Acknowledgments

We thank our resident entomologist, Dr. Jason Chen, for his help in navigating the biology of this system. We are grateful to Speedy Worm for their supplying these larvae. Funds for this research were provided by a grant from the US National Institute of General Medical Sciences via R35GM136407. Roland R. Regoes was supported by the Swiss National Science Foundation (grant 31003A_179170). This work was supported by the Emory University Integrated Cellular Imaging Core Facility (RRID:SCR_023534). The content of this article is solely the responsibility of the authors and does not necessarily reflect the official views of the National Institute of Health or the Swiss National Science Foundation. The funders had no role in the design or execution of this research nor the decision to publish the results.

