## Supplementary material for "The role of innate immunity, antibiotics, and bacteriophages in the course of bacterial infections and their treatment": All Supplemental Materials

Corresponding Author: Bruce R. Levin

**This PDF file includes:**

Supporting text

Tables S1

Figures S1 to S7

SI References

Text S1. Model Equations and Description

To investigate the within-host dynamics of bacterial infections in the experimental *Galleria mellonella* system, we developed a mathematical population model. The model is described by the following system of ordinary differential equations:

$$\frac{dU}{dt}=\left( 1-\frac{U}{K_{U}} \right)rU-h_{1}EU-f\left( 1-\frac{P}{K_{P}} \right)U+b\left( 1-\frac{U}{K_{U}} \right)P$$

$$\frac{dP}{dT}=\left( 1-\frac{P}{K_{P}} \right)rP+f\left( 1-\frac{P}{K_{P}} \right)U-b\left( 1-\frac{U}{K_{U}} \right)P$$

$$\frac{dE}{dt}=\left( a+sU \right)\left( E_{tot}-E-\frac{h_{2}}{g}EU \right)-dE$$

We assume that bacteria can be present in both an unprotected compartment ($U$), where they are exposed to the immune effectors ($E$), and a protected compartment ($P$), where bacteria are shielded from the immune effectors. Bacteria in both compartments, $U$ and $P$, grow exponentially at rate $r$, with their growth limited by the carrying capacities $K_{U}$and $K_{P}$, respectively, where $K_{P}<K_{U}$. Bacteria migrate back and forth between the unprotected and the protected compartment at maximal rates $f$ ($U$ to $P$) and $b$ ($P$ to $U$). These rates are proportionally reduced as the destination compartment approaches carrying capacity.

In our model the innate immune system is described by a population of activated immune effectors ($E$). Following Pilyugin and Antia (2000)(1) we assume a fixed total number of immune effectors ($E_{tot}$). Un-activated immune effectors are described by the term $(E_{tot}-E-\frac{h_{2}}{g}EU)$, where $\frac{h_{2}}{g}EU$ represents the number of activated effectors currently engaged with unprotected bacteria ($U$, details below). The un-activated effectors activate with a background rate $a$ or in response to bacteria as described by the term $sU$. Activated effectors ($E$) deactivate with rate $d$.

Mirroring the natural mode of action of immune systems, where pathogens are incapacitated upon contact with activated immune effectors, unprotected bacteria $U$ are removed by $E$ at rate $h_{1}$ according to mass action kinetics. Specifically, we assume that this interaction between $E$ and $U$ is a two-step process. First, $E$ encounter $U$ via mass-action kinetics by rate $h_{2}$ – this is analogous to how phagocytes capture bacterial cells. Then, they kill them at rate $g$ – mirroring the enzymatic degradation that occurs within phagocytes. Therefore, the ratio $\frac{h_{1}}{h_{2}}$describes how many bacteria an effector can simultaneously phagocytize.

Assuming that killing is relatively fast (i.e. $g$ is large) allows us to make a quasi-steady state assumption for the dynamics of the engaged immune effectors and integrate the handling time via $\frac{h_{2}}{g}EU$into the available number of activatable immune effectors (see (1) for details). Intuitively, the duration of killing by immune effectors results in a reduction of available effectors that could kill bacteria at a rate of $\frac{h_{2}}{g}EU$.

Table S1. Model parameter values.

| **Parameter** | **Description** | **Value** | **Range** | **Sampling** |
| --- | --- | --- | --- | --- |
| inoculum | Starting population size of bacteria | 10 – 10^7.5^ | 10 – 10^7.5^ | Log uni |
| $r$ | Maximal growth rate in absence of antibiotics | 1 | 1 | / |
| $K_{U}$ | Carrying capacity in protected site | 10^8^ | 10^8^ | / |
| $K_{P}$ | Carrying capacity in unprotected site | 1500 | 10^2^ – 10^4^ | Log uni |
| $f$ | Migration rate to protected site | 0.001 | 10^-7^ – 0.1 | Log uni |
| $b$ | Migration rate to unprotected site | 0.1 | 10^-7^ – 0.1 | Log uni |
| $E_{0}$ | Starting population size of activated immune effectors | 1000 | 1 – 10^5^ | Log uni |
| $E_{tot}$ | Total number of available immune effectors (2, 3) | 10^5^ | 10^4.5^– 10^5.3^ | Log uni |
| $h_{1}$ | Killing rate of bacteria by immune effectors | 0.001 | 10^-7^ – 0.1 | Log uni |
| $h_{2}$ | Engagement rate of activated immune effectors with unprotected bacteria | 0.001 | 10^-7^ – 0.1 | Log uni |
| $d$ | Deactivation rate of immune effectors | 0.01 | 10^-4^ – 1 | Log uni |
| $a$ | Background activation rate of immune effectors | 0.01 | 10^-4^ – 1 | Log uni |
| $g$ | Handling rate of immune effectors | 0.5 | 10^-3^ – 4 | Log uni |
| $s$ | Activation rate of immune effectors in response to unprotected bacteria | 0.001 | 10^-7^ – 0.1 | Log uni |

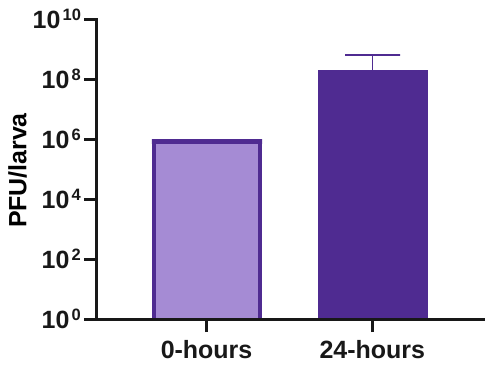

Fig. S1. Phage density before and after treatment. Larvae were infected with 10^8^ CFU/larva of *S. aureus* MN8 and immediately treated with 10^6^ PFU/larva of PYO^Sa^. After 24 hours, larva were sacrificed and the phage density determined (N=20 larvae). Shown are means and standard deviations. The change in the phage density pre- and post-infection is highly statistically significant (*p*<.0001 ****).

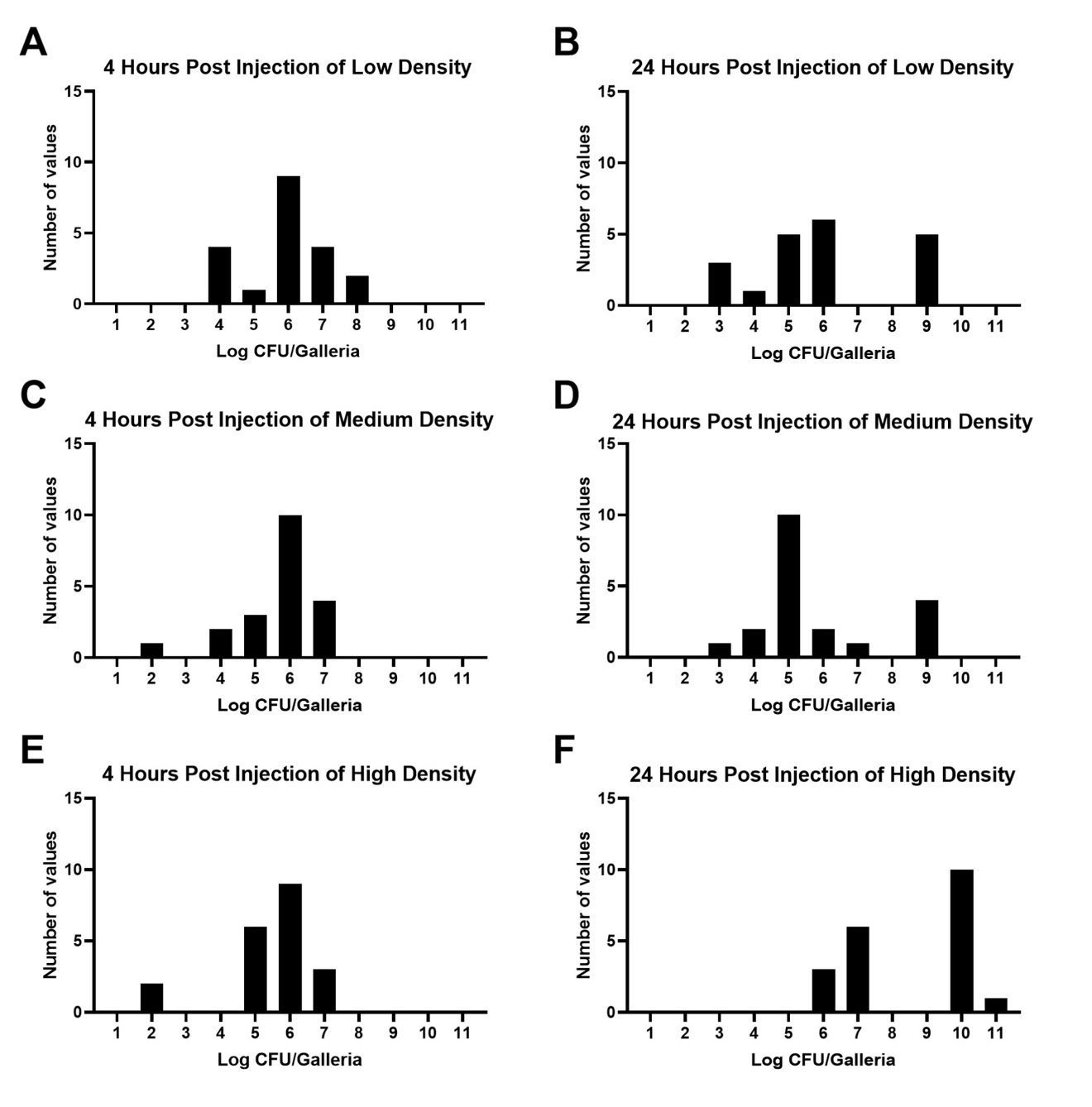

Fig. S2. Dynamics of infection. Larva were infected with three different densities of *S. aureus* MN8: low (10^4^ CFU/larva; A, B), medium (10^6^ CFU/larva; C, D), or high (10^8^ CFU/larva; E, F) and were sacrificed after 4 (A, C, E) or 24 (B, D, F) hours post infection. Shown are the number of larvae within each log_10_ CFU. For each condition N=80.

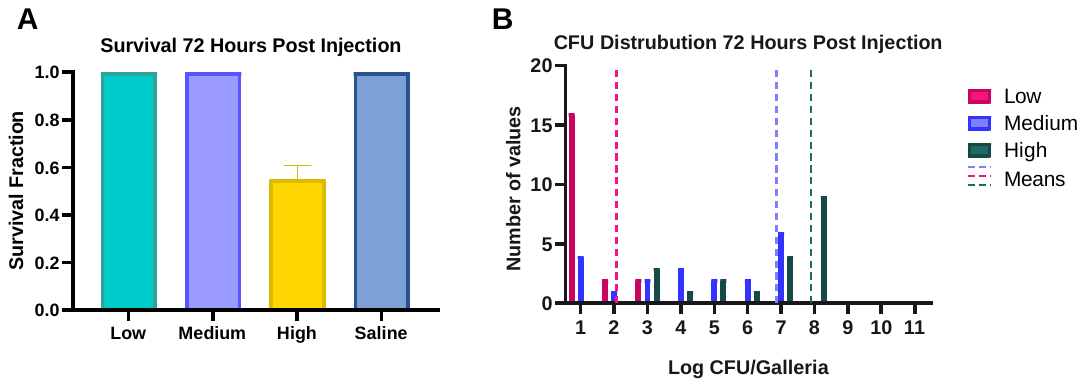

**Fig. S3. Larvae survival and infection dynamics at 72 hours with different bacteria inoculum densities.** (A) Larvae were injected with *S. aureus* MN8 at 10^4^ CFU/larva (turquoise), 10^6^ CFU/larva (purple), 10^8^ CFU/larva (yellow), or saline (blue). Mortality was assessed at 72 hours. Presented are means and standard deviations for the fraction of survival (N= 20 larvae). Low, medium, and high differ with saline at *p*<.0001 ****; low is statistically different from high at *p*<.001 ***; medium is different with high at *p*<.001 ***; and low and medium are not statistically different. (B) Larva were infected with three different densities of *S. aureus* MN8: low (10^4^ CFU/larva; pink), medium (10^6^ CFU/larva; purple), or high (10^8^ CFU/larva; green) and were sacrificed after 72 hours post infection. Shown are the number of larvae within each log_10_ CFU. The mean of the CFU at each timepoint for each inoculum density is presented as a dotted line of the same color. For each condition N=20 individual larvae.

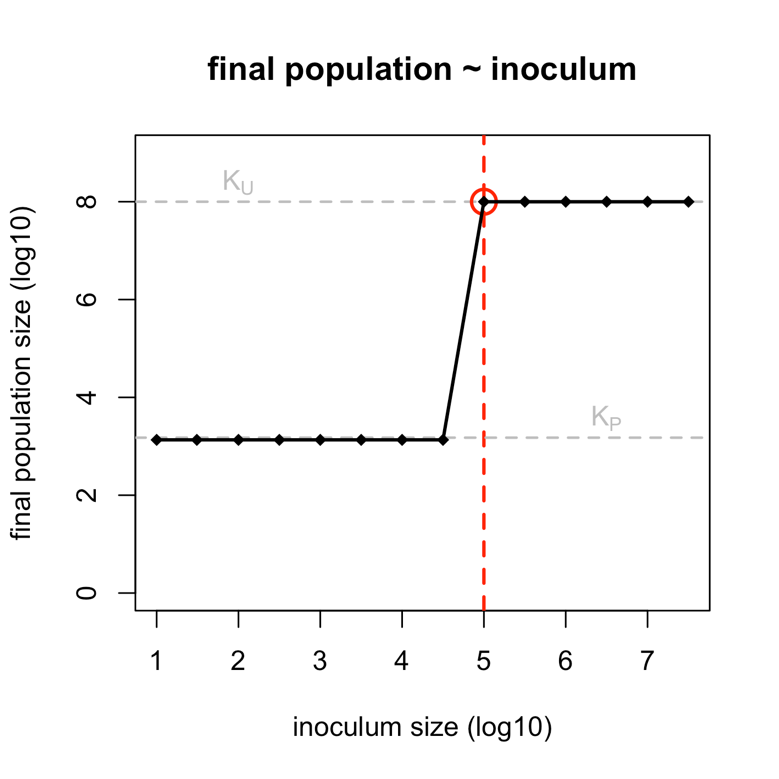

**Fig. S4. High bacterial inoculum sizes overwhelm the immune system.** Shown are the simulated total bacterial population sizes ($U + P$) at 72 hours after inoculation for different inoculum sizes ($U_{0}$). Grey, dashed lines show the carrying capacities of the unprotected ($K_{U}$) and the protected site ($K_{P}$). The critical inoculum threshold– the first size of inoculum that overwhelms the immune system – is highlighted in red. See Table S1 for parameter values used.

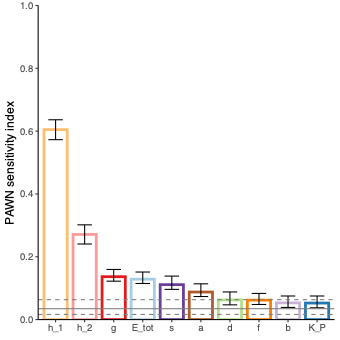

**Fig. S5. Influence of model parameters on the critical inoculum threshold.** We ran simulations for 10,000 randomly generated parameter sets to determine the critical inoculum thresholds – the first inoculum which overwhelms the immune system (highlighted in red in Fig. S4). For 1479 parameter sets, the immune system was not overwhelmed by any tested inoculum sizes. The remaining 8,521 parameter sets showed a clear critical inoculum threshold. For these 8,521 threshold values, we conducted a global sensitivity analysis by calculating PAWN sensitivity indices with bootstrapped 95% confidence intervals (CIs) to assess which model parameters most influence the threshold. The solid and dashed grey lines show the sensitivity index and CI of a dummy variable, i.e., a parameter which has no influence on the model output. The two immune parameters, h1 and h_2_, are indicated to have the most influence on the critical inoculum threshold, whereas the non-immune parameters, i.e., 𝑏, 𝐾P, and 𝑓, the least. See Material and Methods for details and Table S1 for parameter ranges sampled.

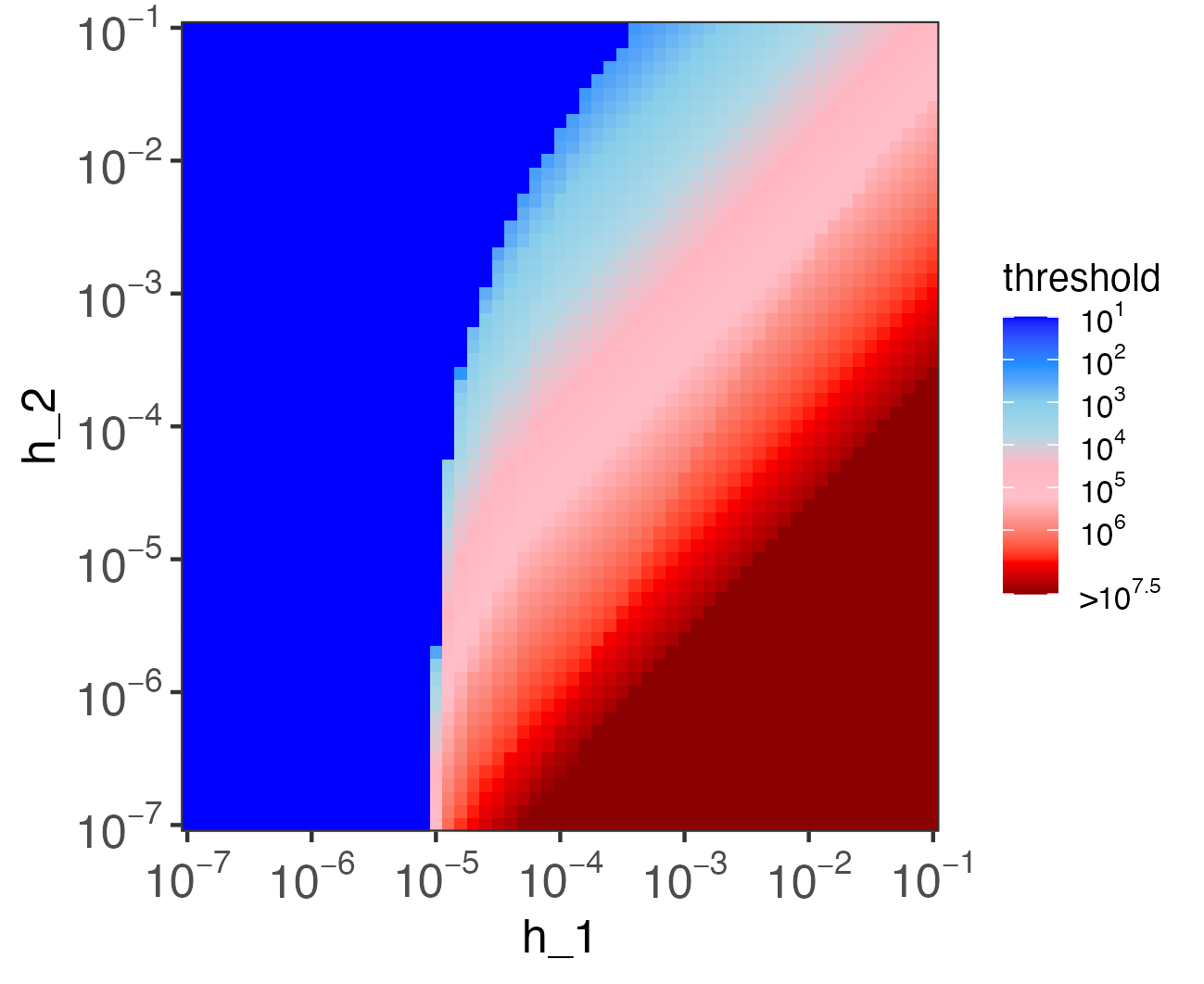

**Fig. S6. Influence of the immune parameters,** $\boldsymbol{h}_{\boldsymbol{1}}$ **and** $\boldsymbol{h}_{\boldsymbol{2}}$**, on the critical inoculum threshold.** Shown is a heatmap of simulated inoculum thresholds – from low (blue) to high (red) – for pairwise combinations of effector killing rate ($h_{1}$) and effector engagement rate ($h_{2}$). All other model parameters are kept constant. For low values of $h_{1}$ (<10^-5^) the immune system is overwhelmed with minimal bacterial inocula – irrespective of $h_{2}$. When $h_{1}$>10^5^ and $h_{2}$is at low to intermediate values of (<10^-4^) the critical threshold increases to the point where no tested inocula can overwhelm the immune system (i.e. threshold >10^7.5^). At high values of $h_{1}$ and $h_{2}$ the threshold is intermediate, ranging from 10^3^-10^5^. See Table S1 for parameter values and Fig S4 for determination of the critical inoculum threshold.

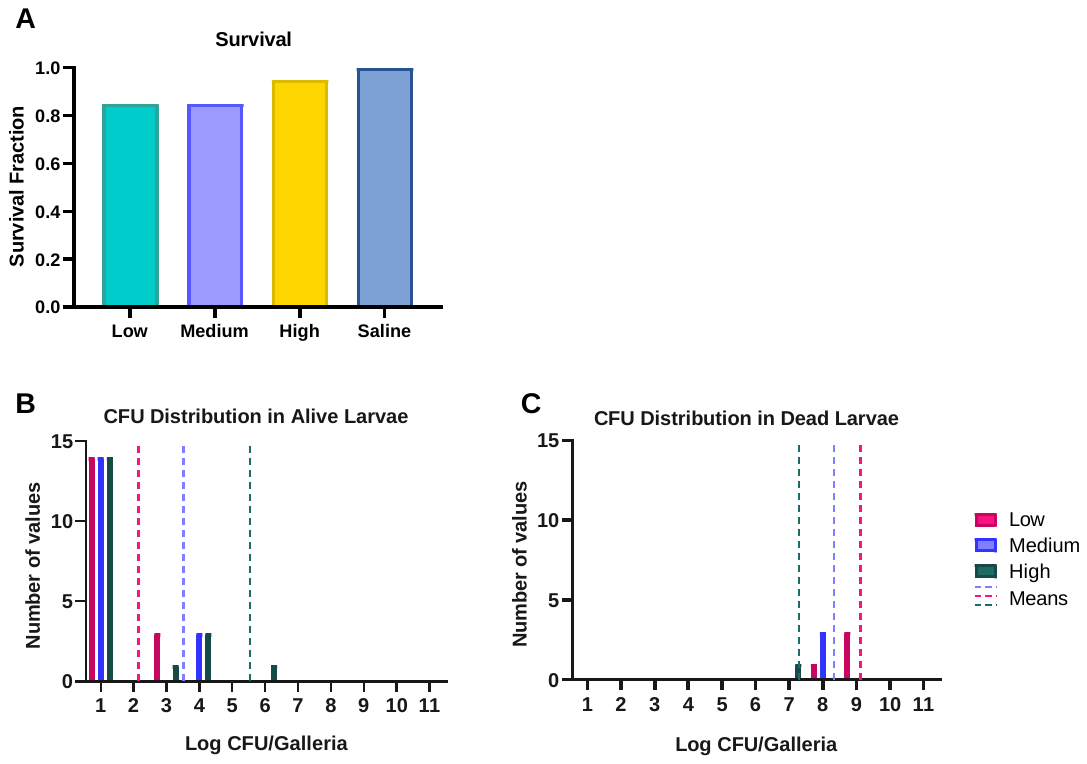

**Fig. S7. Larvae survival and infection dynamics at 24 hours with different bacteria inoculum densities and ablated innate immune system.** (A) Larvae were injected with *S. aureus* MN8 at 10^4^ CFU/larva (turquoise), 10^6^ CFU/larva (purple), 10^8^ CFU/larva (yellow), or saline (blue). Mortality was assessed at 24 hours. Presented are means and standard deviations for the fraction of survival (N= 20 larvae). Low, medium, and high survival fractions are not statistically different. (B and C) Larva were infected with three different densities of *S. aureus* MN8: low (10^4^ CFU/larva; pink), medium (10^6^ CFU/larva; purple), or high (10^8^ CFU/larva; green) and were sacrificed after 24 hours post infection. Shown are the number of larvae within each log_10_ CFU. The mean of the CFU at each timepoint for each inoculum density is presented as a dotted line of the same color. For each condition N=20 individual larvae. (B) Larvae that were alive at 24 hours. (C) Larvae that were dead at 24 hours.
